# A biologically plausible dynamic deep network for recognizing structure from motion and biological motion

**DOI:** 10.1101/2022.08.18.504369

**Authors:** Anila Gundavarapu, V Srinivasa Chakravarthy

## Abstract

A breakthrough in the understanding of dynamic 3D shape recognition was the discovery that our visual system can extract 3D shape from inputs having only sparse motion cues such as (i) point light displays and (ii) random dot displays representing rotating 3D shapes - phenomena named as biological motion (BM) processing and structure from motion (SFM) respectively. Previous psychological and computational modeling studies viewed these two as separate phenomena and could not fully identify the shared visual processing mechanisms underlying the two phenomena. Using a series of simulation studies, we describe the operations of a dynamic deep network model to explain the mechanisms underlying both SFM and BM processing. In simulation-1, the proposed Structure from Motion Network (SFMNW) is trained using displays of 5 rotating surfaces (cylinder, cone, ellipsoid, sphere and helix) and tested on its shape recognition performance under a variety of conditions: (i) varying dot density, (ii) eliminating local feature stability by introducing a finite dot lifetime, (iii) orienting shapes, (iv) occluding boundaries and intrinsic surfaces (v) embedding shape in static and dynamic noise backgrounds. Our results indicate that smaller dot density of rotating shape, oriented shapes, occluding boundaries, and dynamic noise backgrounds reduced the model’s performance whereas eliminating local feature stability, occluding intrinsic boundaries, and static noise backgrounds had little effect on shape recognition, suggesting that the motion of high curvature regions like shape boundaries provide strong cues in shape recognition. In simulation-2, the proposed Biological Motion Network (BMNW) is trained using 6 point-light actions (crawl, cycle, walk, jump, wave, and salute) and tested its action recognition performance on various conditions: (i) inverted (ii) scrambled (iii) tilted (iv) masked (v) actions, embedded in static and dynamic noise backgrounds. Model performance dropped significantly for the presentation of inverted and tilted actions. On the other hand, better accuracy was attained in distinguishing scrambled, masked actions, performed under static and dynamic noise backgrounds, suggesting that critical joint movements and their movement pattern generated in the course of action (actor configuration) play a key role in action recognition performance. We also presented the above two models with mixed stimuli (a point light actions embedded in rotating shapes), and achieved significantly high accuracies. Based on the above results we hypothesize that visual motion circuitry supporting robust SFM processing is also involved in the BM processing. The proposed models provide new insights into the relationships between the two visual motion phenomena *viz.*, SFM and BM processing.

## 1. INTRODUCTION

The computational principles underlying extraction of structure from motion (SFM) and perception of human/biological motion are still not fully understood. A 1953 study by Wallach and O’Connell (Wallach & O’connell, 1953), on structure from motion, tested the kinetic depth effect and concluded that an unfamiliar 3D object can be recovered from the orthographic projection of changing shadow, provided the object producing the shadow is rigid. On similar lines, several other studies demonstrated that the three-dimensional (3D) structure of an object can be recovered from two-dimensional projections of transparent, revolving cylinders with dots on its surface (Inada et al., 1987; Ramachandran, 1985; Ramachandran et al., 1987; Schwartz & Sperling, 1983; Ullman, 1979, 1983).

Studies have shown that middle temporal area (MT), an area that lies on the dorsal (“where”) pathway of the primate visual system, is directly involved in SFM perception (Bradley et al., 1998). Lesion studies showed that monkeys with MT lesions display impairment in tasks where three-dimensional structure is to be perceived from motion cues alone (Siegel & Andersen, 1988, 1990). A study that used bistable SFM stimulus (Grunewald et al., 2002), tested the effect of SFM stimuli on neural activity in cortical areas V1 and MT, and reported that MT is directly involved in the generation of SFM percept, whereas V1 activities are indirectly related to SFM perception which is due to top-down feedback from MT. However, given its complexity, SFM involves a high-level of visual processing. While perceiving shape in a given rotating stimulus the movements of individual dots can be encoded by low- and mid-level motion processes, but inference of 3D shape from variations in 2D motion requires sophisticated high-level process, operating over large areas of display.

The original kinetic depth experiments were performed using wireframe objects. On the other hand, later studies have established that structure can also be perceived from displays of isolated points. An early study (Johansson, 1973) demonstrated our ability to perceive human form walking/dancing, simply by observing moving point-lights fixed at the major joints of a subject’s skeleton moving in dark, commonly referred to as biological motion displays or point-light (PL) displays. Subsequently, others have uncovered a wide range of abilities related to the perception of PL displays. For example, subjects are able to determine gender (Mather & Murdoch, 1994), emotional state (Atkinson et al., 2004, 2007), age (Montepare & Zebrowitz-McArthur, 1988), and identity of a person (Troje et al., 2005) based on the information provided by PL displays. Moreover, perception of human action remains robust even when PLs are embedded in a noisy background (Cutting et al., 1988). Furthermore, biological motion perception has been demonstrated in vertebrates such as mice (Atsumi et al., 2018), cats (Blake, 1993), dogs (Delanoeije et al., 2020; Eatherington et al., 2019), chicks (Vallortigara et al., 2005; Vallortigara & Regolin, 2006) and pigeons (Dittrich et al., 1998); also, in invertebrates like spiders (de Agrò et al., 2021); and in 4-month-old human infants (Bertenthal, 1993). The evidence suggests that the mechanisms involved in biological motion perception become active very early in an organism’s development and serve as a kind of “life detectors” (Troje & Westhoff, 2006).

A class of findings on biological motion support the hypothesis that the form and motion cues together play a role in reliable perception of biological movement (Lange et al., 2006; Lange & Lappe, 2007; Pinto & Shiffrar, 1999). Functional imaging studies (Grossman et al., 2000), as well as single-cell recordings (Oram & Perrett, 1994) indicate the existence of specific mechanisms for the processing of biological motion within the area Posterior Superior Temporal Sulcus (STSp). Lesion studies and neuroimaging studies have implicated that the STSp is critical to biological motion perception, while lower-level motion processing takes place in MT (Beauchamp et al., 2002; Bonda et al., 1996; Boussaoud et al., 1990; Felleman & van Essen, 1991; Grossman et al., 2000; Saygin, 2007; Servos et al., 2002; Vaina et al., 2001; Vaina & Gross, 2004). Furthermore, it has been suggested that STSp is an area where motion and form information, coming from dorsal and ventral visual streams, are integrated (Blake & Shiffrar, 2007). Studies have also shown that performance in perceiving biological motion is impaired substantially when an inverted point-light actor is displayed (Bertenthal & Pinto, 1994; Pinto & Shiffrar, 1999; Sumi, 1984).

On the other hand, theoretical studies have reported that a unique structure can, in principle, be recovered from motion information alone, which is integrated over a small extent in space and time (Barron, 1984; Hildreth & Koch, 1987; Ullman, 1983). However, with regard to the human visual system, the structure can be perceived either due to the spatial extent of viewed motion, or temporal extent of viewed motion or both. 2D projections of rotating transparent objects can be considered as examples of the spatial extent of the viewed motion and those of biological point light displays can be considered as the temporal extent of viewed motion. We present a a computational model, that consists of two stages analogous to V1-MT, and can recognize both SFM and biological motion.

In this paper we describe a dynamic deep neural network model for finding structure from motion and biological motion. We test the proposed model with both rotating shapes and biological point-light displays, and drawn conclusions based on the network performance. Earlier we presented a biologically plausible computational model (Gundavarapu et al., 2019) that can explain neural responses at V1 and MT to motion stimuli. This dynamic model was recently extended using a deep convolutional neural network (CNNs) for optic flow recognition and was described in a companion paper (Gundavarapu and Chakravarthy, 2022). On similar lines, in the present paper, we describe two models viz., Structure from Motion Network (SFMNW) and Biological Motion Network (BMNW) designed to model higher order perceptual phenomena like structure from motion and biological motion respectively.

We perform a series of 5 simulations to test the two models. In simulation-1, we trained the SFMNW (structure from motion network) with rotating shapes and tested its generalization performance on oriented, masked, varied density and varied dot lifetime displays. In simulation-2, we trained BMNW (biological motion network) using 6 point-light action sequences to discriminate actions from point-light displays of different conditions such as inverted, scrambled, oriented and masked inputs. We then investigated the behavior of SFMNW and BMNW on mixed stimuli: point-light actor enclosed by a rotating transparent shape (cone, cylinder, sphere) and studied the network performance in action recognition and shape recognition in simulation-3. In simulations 4 and 5 we tested whether both the models SFMNW, BMNW can recognize shapes and actions embedded in different types of static and dynamic noisy backgrounds.

In summary, our work makes the following major contributions:

a. proposed a computational framework to simulate structure from motion and biological motion
b. demonstrate successful model performance under a wide range of display formats such as orientation, inversion, scrambled, static and dynamic noise, dot density and lifetime and masking etc.

## 2. ARCHITECTURE PROPOSED

SFMNW and BMNW are designed using a dynamic deep network proposed below. SFMNW is designed and trained to recognize the 3D shape from rotating 2D stimuli, so that the neurons learn to extract a 3D shape from abstract local motion. On the other hand, BMNW is trained to recognize the actions from point light displays, so that the neurons here learn to recognize the human form of action from sparse local motion cues.

The proposed dynamic deep network (Figure. 1) consists of a cascade of two stages: 1) Velocity selective mosaic network (VSMN), and 2) Convolutional neural network (CNN). The two stages are trained successively, VSMN first, followed by CNN.

**Figure 1:**
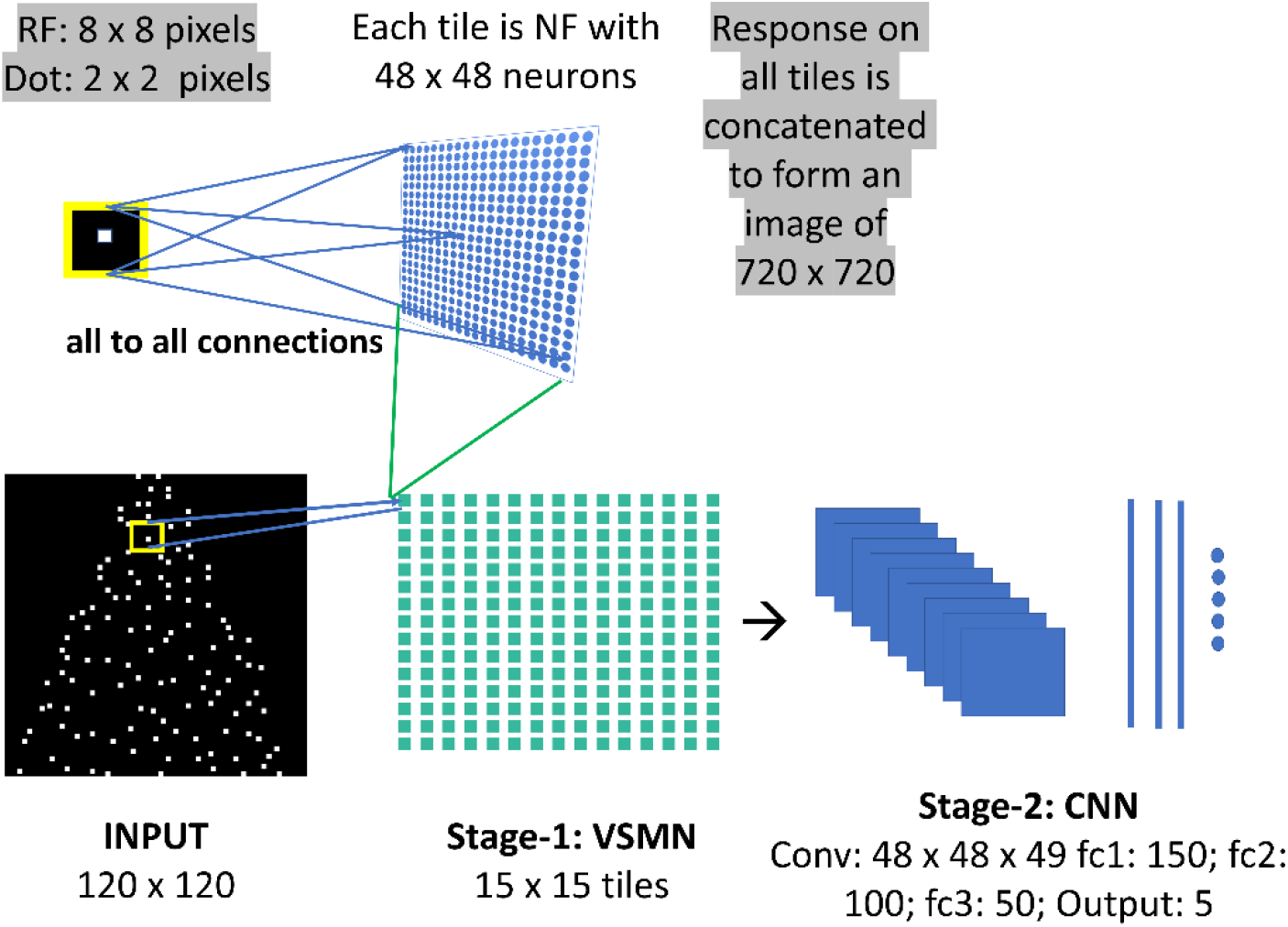
Dynamic deep neural network architecture: The network consists of two stages: (1) Velocity selective mosaic network (VSNW) where neurons encode direction and speed of a dot present with in the receptive field (RF), (2) Convolutional neural network (CNN) that takes VSMN response as input and learn representations to categorize 3D shapes or human actions. This figure depicts the SFMNW1 used in simulation-1. We present two more networks: SFMNW2 and BMNW; each follows the same architecture with variation in number of fully connected layers in CNN as described in Table 2.

### 2.1 Velocity selective mosaic network (VSMN)

VSMN consists of a 15 × 15 “mosaic” in which each “tile” is a 2D array of neurons (Fig. 1). A single tile is an independent neural field (NF) network wherein the neurons are connected among themselves by lateral connections. In a given NF, each neuron receives inputs from a shared patch of size 8 × 8 of the input image. Thus, all the neurons in a given NF have a shared receptive field of size 8 × 8. The input image of size 120 × 120 can be considered as being composed of 15 × 15 array of small patches each of size 8 × 8. Even though all neurons in a given NF respond to the same patch of the input image, they develop preferences to different motion directions and speeds due to the initial variation in the afferent connections, and the competition created by the lateral connections. As a result of training, NF neurons were clustered into populations in such a way that each population is selective to a specific direction and speed of motion of a dot.

### 2.2 NF training procedure

Each NF in VSMN is composed of 48 × 48 neurons. All these neurons receive common input from 8 × 8 window of the input image of size 120 × 120. Despite the common input received by all NF neurons, their initial response varies according to the random afferent weights. The net afferent input of NF neuron *(i, j)* is calculated as,

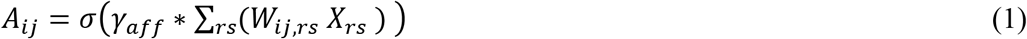

where *X* is 8 × 8 image window which constitutes the Receptive Field (RF) of the neuron at *(i, j)*. *W* is afferent weight matrix of neuron at *(i, j)*. *W_ij,rs_* is the afferent weight connection from pixel position *(r, s)* to *(i, j)*. *γ_aff_* is a constant scaling factor and is initialized before training begins. *σ* is piecewise linear sigmoid activation function described in the Methods section.

The initial response is dominated by afferent input which is further modified by the lateral interactions. Each neuron *(i, j)* in NF maintains two types of lateral connections (Fig. 2B): i) short-range excitatory lateral connections with the neurons within a neighbourhood of radius *r_exc_*. ii) Long-rage inhibitory lateral connections with the neurons within a neighbourhood of radius *r_inhb_* and outside the radius *r_exe_*. In other words, two neurons in a NF have an inhibitory lateral connection if their mutual distance is more than *r_exc_* and less than *r_inhb_*. The *r_exc_* and *r_inhb_* values are uniform for all the neurons within and across the tiles.

**Figure 2:**
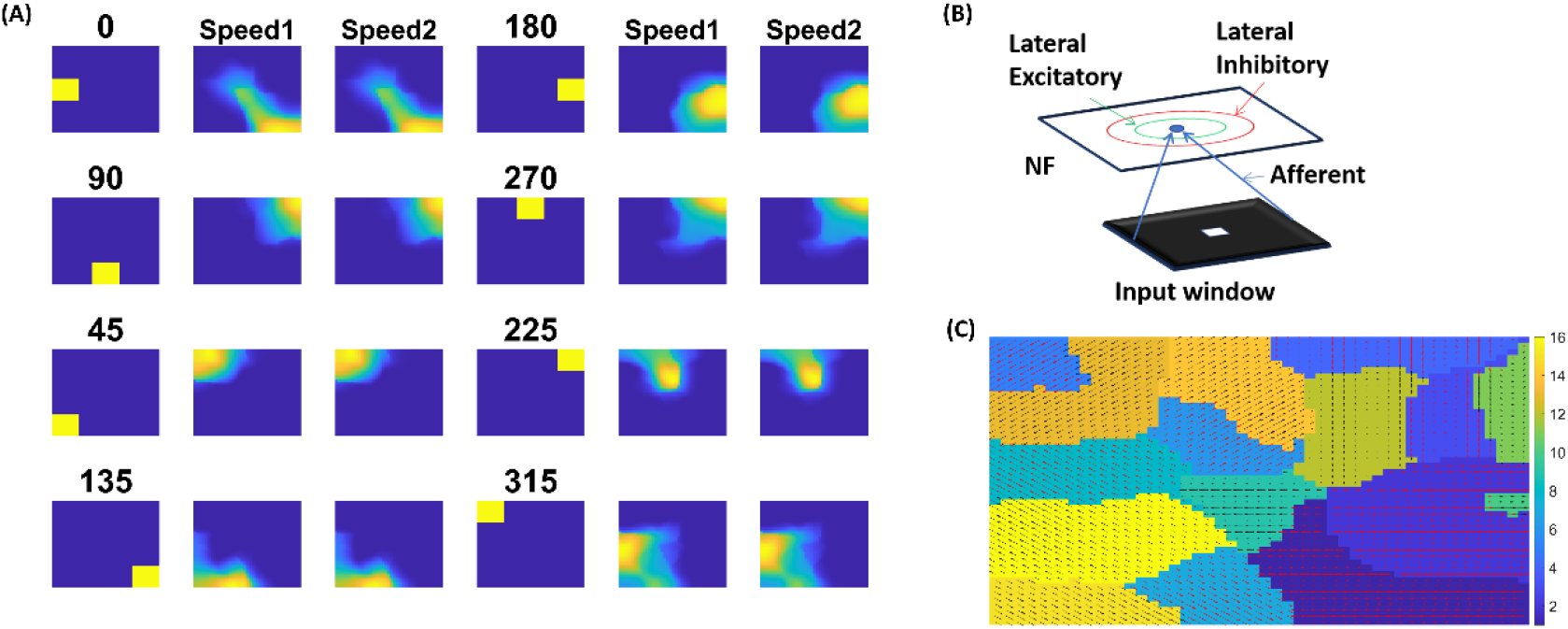
Neural field network (NF) that encode speed along with direction: In (A) 1^st^ and 4^th^ columns display the first frame (8 × 8) of an input sequence that consists of a single dot (2 × 2) moving in a specific direction at two different speeds. Whereas the 2^nd^ and 5^th^ columns show NF responses (48 × 48) at speed-1, the 3^rd^ and 6^th^ columns show responses at speed-2. (B) shows NF architecture that consists of a 2D array of neurons. Red and green circles indicate excitatory and inhibitory radii and blue lines indicate the afferent connections with the pixels on the input frame. (C) shows the velocity selective map indicating that, even though the same neuron population become selective for the inputs with same direction and different speeds, within the population neurons have different speed preferences.

For several discrete time steps ‘*s’* (settling time), the response of the neuron *(i, j)* is modified by afferent and lateral interactions that takes place simultaneously.

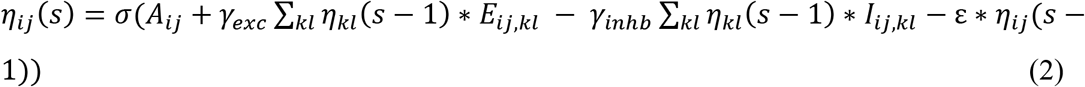

where *η_ij_* stands for the activity of the neuron *(i, j)*, *E_ij,kl_* and *I_ij,kl_* are excitatory and inhibitory weight connections from the neuron *(k, l)* to *(i, j)* that are randomly initialized before training begins. The relative strengths of excitatory and inhibitory lateral effect are controlled by the constant scaling factors *γ_exc_* and *γ_inhb_*. The above dynamical equation can recognize variation in speed along with the direction of motion, i.e., velocity. During our simulations we tried various scaling values (*ε*) for *η*_*ij*_ (*s* − 1) ranging from 0.1 to 0.001. At higher values of *ε*, the network fails to distinguish speed. At lower values, the network response is unstable during the presentation of sequence. In other wards lateral interactions could not produce unique activity pattern in NF to encode speed feature. The value we chose for *ε* is 0.01.

At the end of *‘s’* time steps, NF response settles down and all the three types of weights (afferents, excitatory laterals and inhibitory laterals) are updated. Weights are updated for presentation of each image in the input sequence. Let ‘*t*’ represents the time of presentation of current frame to NF with assumption that at *t=1*, frame-1 is presented. The afferent connections are adapted using symmetric Hebbian rule as specified in eqn. (3) below.

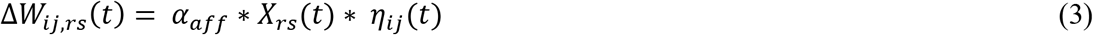

where *α*_*aff*_ is learning parameter for afferent weight connection. *X* is 8 × 8 image window from which neuron *(i, j)* receives input. *W_ij,rs_* is afferent weight connection between the pixel position *(r, s)* and the neuron *(i, j)*. *η_ij_* is the activity of neuron *(i, j)* after settling process.

The lateral connections are adapted using asymmetric Hebbian learning rule as specified in eqn. (4) below.

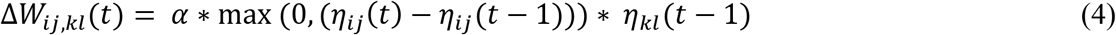

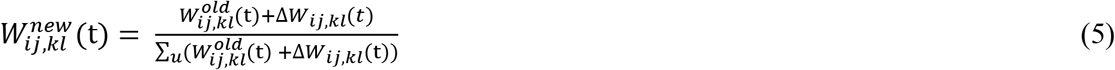

where *ɳ_ij_(t)* is the activity on the neuron *(i, j)* produced in response to the current frame (the frame presented at time ‘*t*’), *ɳ_ij_(t-1)* is the activity on the neuron *(i,j)* for the previous frame. *α* is the learning parameter that controls the rate of learning. Separate learning parameters were used for both excitatory (*α*_*exc*_) and inhibitory (*α*_*inhb*_) connections. All the three types of weight connections are normalized separately as shown in the eqn. (5) to prevent neuron activity growing out of bounds. Various parameters used in the simulation were described in Table 1.

**Table 1:** Parameters used to train NF in VSMN

| Parameter | Tile in VSMN |
| --- | --- |
| NF Dimension | 48 x 48 |
| Receptive field | 8 x 8 |
| $r_{exc}$ | 2 |
| $r_{inhb}$ | 4 |
| $\gamma_{aff}$ | 1 |
| $\gamma_{exc}$ | 50 |
| $\gamma_{inhb}$ | 1.5 |
| $\alpha_{aff}$ | 0.05 |
| $\alpha_{exc}$ | 0.05 |
| $\alpha_{inhb}$ | 0.05 |
| Time step (s) | 10 |
| Epochs | 250 |
| Image size | 80 x 80 |

Note that the training procedure described above is for one tile (a NF) that takes input from a small window (8 × 8) of input image. First, one tile/NF is trained using input sequences made up of 2 × 2 tiny squares (the “dots”) moving in 8 different directions and in each direction at 2 different speeds. Thus, the training set of NFs in VSMN consists of 16 sequences. The response of trained NF, and the direction preferences of NF population are plotted in Fig. 2A and 2C respectively. The weights of a trained tile are copied to the remaining tiles, since different tiles perform identical computations but on different parts of the image.

After training, VSMN weights are frozen, and its responses are used to train the CNN, which consists of one convolutional layer followed by 3 fully connected layers (as shown in Fig. 1) and is trained to recognize: (i) shape of a rotating 2G pattern in SFMNW, (ii) the type of action in PL displays. CNN is designed and simulated using MATLAB 2020a deep learning tool box.

## 3. SIMULATION-1: *Discriminating shapes using motion cues and the role of different variables on shape recognition*

It is well-recognized that affine geometrical information related to 3-D shape can be recovered from orthographically projected motion sequences (Ramachandran, 1985; Ramachandran et al., 1987; Ullman, 1979). Although counter-intuitive, these motion sequences contain adequate information to enable perception of three-dimensional (3-D) shape. For the sake of convenience, we shall refer to such potential information as a set of variables (as described below) and conduct various simulations to investigate the role of each variable in shape recognition.

i. the minimal density of dots required to perceive the structure of a smooth curved surface
ii. the role of temporal correspondence of moving points across frames
iii. the role of edges, corners and the orientation at which the shape depicted

To this day, however, it remains unclear exactly which 2-D image properties are detected and exactly which 3-D structural relations are perceived when observers view structure from motion displays.

### 3.1 Stimuli Creation

The 3D surfaces of 5 different shape sequences (Cone, Cylinder, Sphere, Ellipsoid, and Helix) were defined by moving dots and were created by a computer program using MATLAB. Each sequence is made up of 15 frames, with frame size 120 × 120. White dots of size 2 × 2 were placed randomly on black background. Approximately 120 dots for Helix and 180 dots for the remaining shapes (Cone, Cylinder, Ellipsoid and Sphere) were used. Different sets of standard motion sequences were created using the equations described in the Methods section.

#### (a) Applying mask

To understand the role of motion information present at the edges and corners in enabling shape perception, we made cut-out parts of all 5 vertically rotating shapes by applying invisible masks (of sizes 30 × 40 or 40 × 40) at various positions on the display as described below and created 5 mask conditions as shown in Fig. 3.

**Figure 3:**
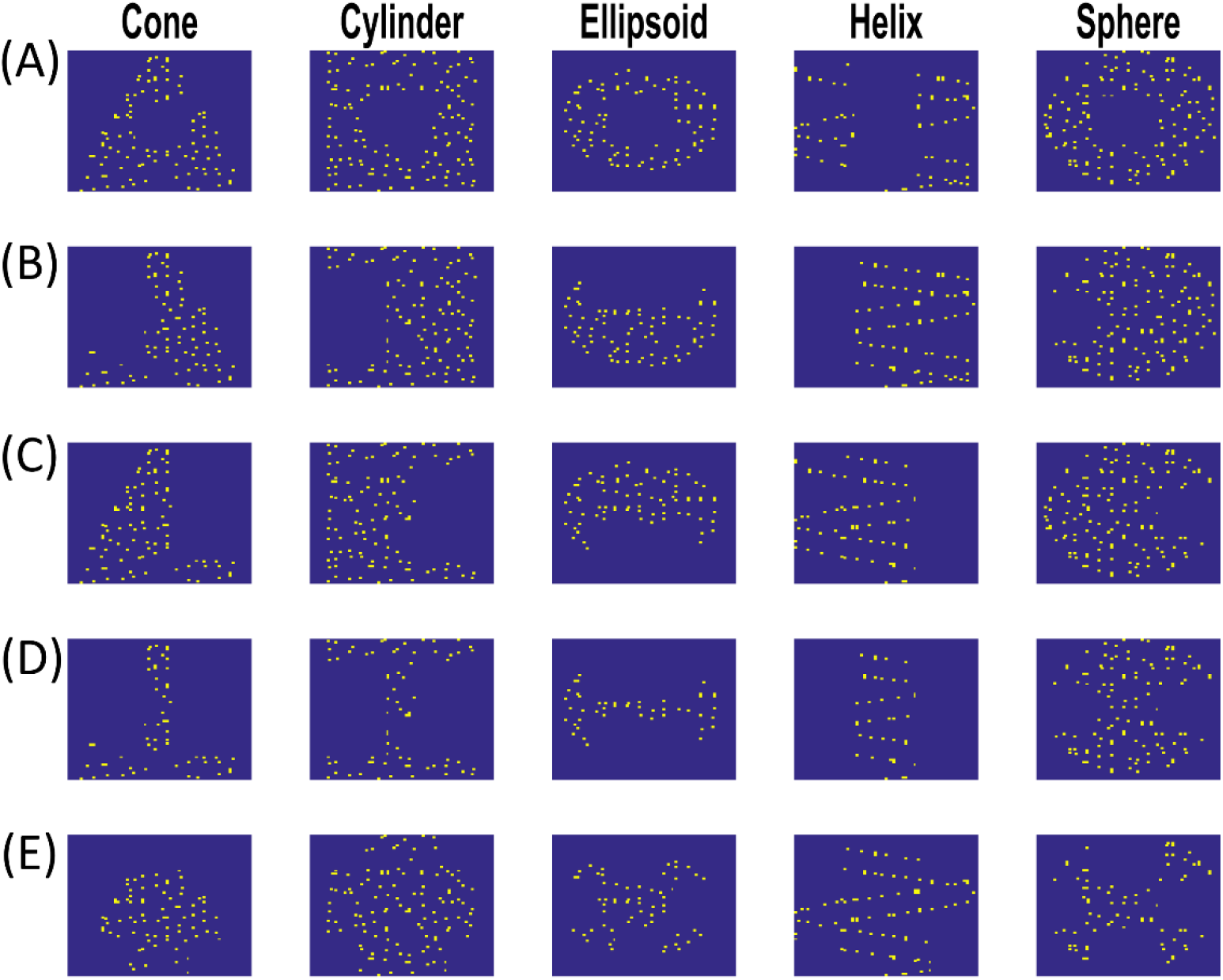
Five different mask conditions used in the simulation: (A) intrinsic surfaces of the shape are occluded. In (B), (C), (D) boundaries of each shape are occluded in 3 different ways. In (E) all corners are occluded to understand the contribution of motion information by boundaries in enabling the 3D shape perception.

i. 40 × 40 size mask centered on of the shape
ii. 40 × 40 size mask on left vertical edge surface (top in case of ellipsoid)
iii. 40 × 40 size mask on right vertical edge surface (bottom in case of ellipsoid)
iv. 40 × 40 sized masks on both the vertical edges together (top and bottom in case of ellipsoid)
v. 30 × 40 sized masks at all corners.

All the above mask positions are stationary and the number of the points in a frame are less in masked stimuli compared to unmasked stimuli. The points on the surface of the shape disappeared when they rotated behind the mask and reappeared on the other side of the mask.

#### (b) Tilted shapes

For each shape class, tilted or oriented 3D shapes were created by tilting its vertical axis of symmetry from 90° to 135° (=90°+45°) and 45° (=90°-45°).

#### (c) Limiting the surface point lifetime

To understand the role of temporal buildup by individual surface points in perceiving SFM, lifetime is assigned to each individual point. The optimal temporal build up is the duration or the number of frames required at which the motion integration through time contributes very little to shape recognition. All the dots present on a surface of a rotating shape are only presented for a predetermined number of frames and are then repositioned. We used point lifetimes of 2, 4, 6, 8 and 15 frames. When the surface point reached the end of its lifetime, it is repositioned to a new random location. Thus, the projected image has fixed dot density. Approximately 25% of dots make short trajectories in each sequence. For example, when lifetime equals 6-time steps, 25% of the surface points appear for 6 frames and are then repositioned randomly with new lifetime. This limits the amount of information contributed by an individual dot to the recovery of the perceived 3-D shape of the object.

#### (d) Varying dot density

To understand the minimum number of dots required for accurate structure perception, different sequences are created by removing various proportions of dots in each sequence, like 25%, 50% and 75% of dots.

### 3.2 Training procedure

A total of 250 rotating random dot sequences (50 for each shape) were created and is divided into training set and test set, each consisting of 175, 75 sequences respectively. The training is performed in two stages. In stage-1, VSMN response is calculated for sequences in training set and test set. In stage-2, the CNN is trained using VSMN responses generated out of training set. Various parameters used to train CNN in SFMNW1 are shown in Table 2. This trained network (SFMNW1) is used here for further analysis and to understand the role of various variables described above. Other sequences like masked, tilted etc., were created using these 250 patterns.

**Table 2:** CNN architecture used to create various networks

| Layer | SFMNW1<br>(Simulation-1) | SFMNW2<br>(Simulation-4) | BMNW |
| --- | --- | --- | --- |
| Input layer | 720 x 720 | 720 x 720 | 720 x 720 |
| Convolution layer (conv1) | 48 x 48 x 49 | 48 x 48 x 49 | 48 x 48 x 49 |
| Fully connected layer (fc1) | 150 x 1 | 1050 x 1 | 150 x 1 |
| Fully connected layer (fc2) | 100 x 1 | 450 x 1 | 50 x 1 |
| Fully connected layer (fc3) | 50 x 1 | 250 x 1 | - |
| Fully connected layer (fc4) | - | 50 x 1 | - |
| Output layer | 5 x 1 | 5 x 1 | 6 x 1 |
| **Learning Parameters** |  |  |  |
| Learning rate | 0.01 | 0.01 | 0.01 |
| Batch size | 5 | 40 | 5 |
| Epochs | 100 | 100 | 100 |

### 3.3 Results

#### Mask

We calculated the accuracy of 3D shape recognition as a function of the mask position and plotted the results in Fig. 4A. Masks are placed on the surface of the rotating shape in 5 different ways to occlude boundaries and intrinsic areas of a shape. When the model is presented with the first mask condition displays (where the intrinsic area of surface shape is partially covered by a mask in Fig. 3A, 3D) shape recognition is not affected. All the shapes are recognized with high accuracy. While presenting mask type 2, 3 displays (where one of the sides of vertically rotating shapes are masked as in Figs. 3B, 3C) the model could recognize all the shapes with high accuracy except the cone. In mask type 4 (where both the sides of the vertically rotating shapes are masked Fig. 3D) performance is degraded in recognizing cone and helix whereas in the case of mask type 5 (where corners of a surface shape are partially covered by a mask as in Fig. 3E) recognition of cone and cylinder are affected. Our results show that the turns in the helix and vertical edges and corners in the cone play important role in their shape recognition.

**Figure 4:**
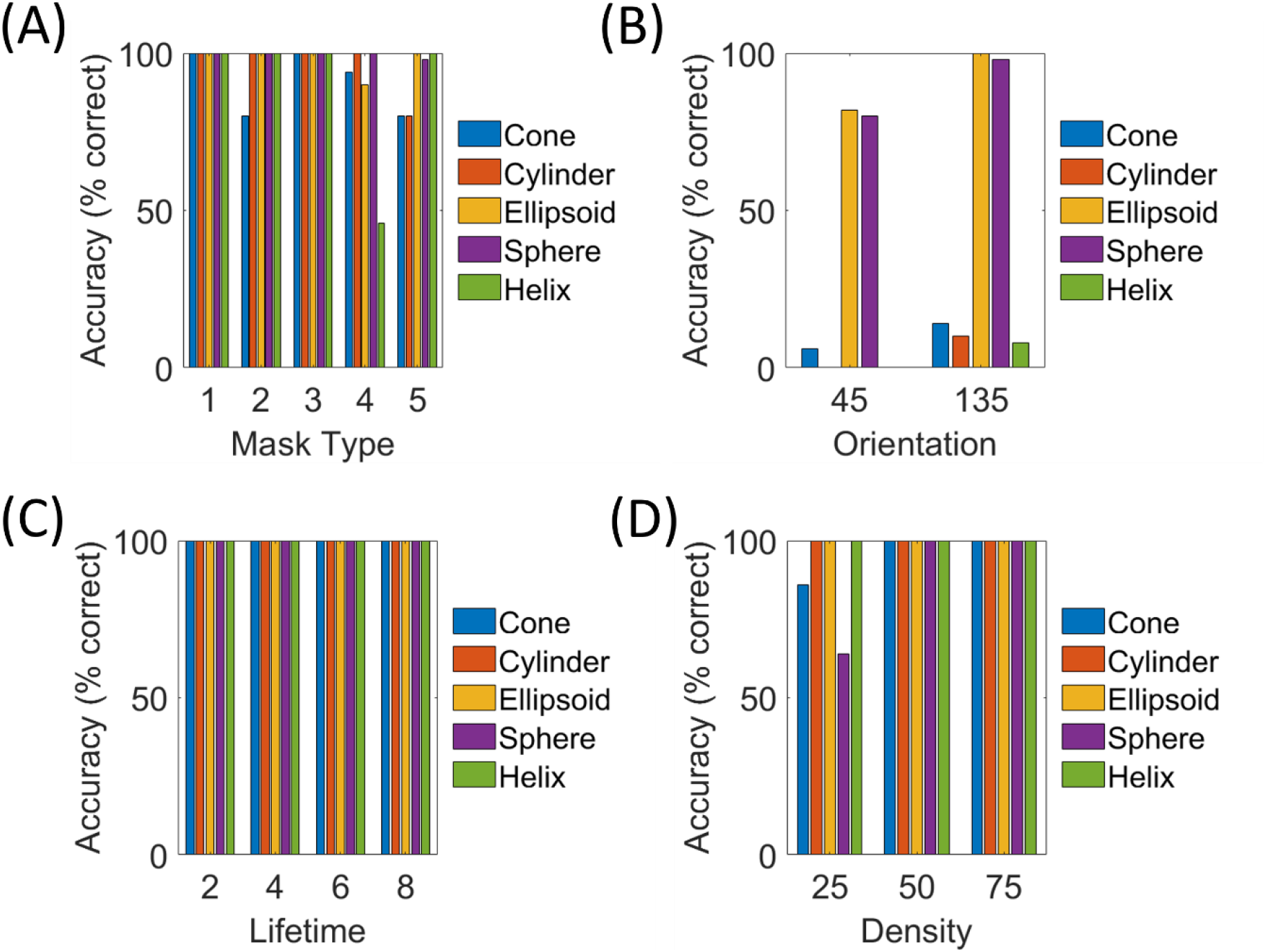
Shape recognition performance of SFMNW1 for various display formats: (A) masked displays, (B) Oriented displays, (C) Lifetime displays, and (D) Varied density displays. Note that all display formats are created using standard displays of training and test sets (i.e., 250 sequences). At a time only one parameter is changed in a sense that while applying mask orientation, lifetime and density is maintained as in as in standard displays and vice versa. Network performance is affected badly in case of oriented shapes and shapes with 25% dot density. On the other hand, for lifetime displays, shapes whose dot density >=50% produced highest accuracy. The network performance is satisfactory for different masked inputs.

#### Orientation

We also tested how well the model that was trained on normal data can recognize the tilted shapes. As shown in Fig. 4B model performance was affected severely. Except ellipsoid and sphere other shapes are not recognized by the model.

#### Assigning lifetime to dots

It is interesting to study the effect of reducing individual image feature stability, by eliminating dots, on shape recognition performance. Individual feature stability can be attenuated by disrupting the temporal correspondences produced by individual points. Here temporal buildup of perceived SFM is manipulated by assigning lifetime to each surface point. Under limited point lifetimes, individual points make only short trajectories, causing an increase in uncorrelated noise. We initially expected that, while the model could recognize 3D shape from motion for longer view point lifetimes (due to longer temporal integration), it suffers disproportionately when the surface point lifetime is reduced. Surprisingly, as shown in Fig. 4C none of the view point lifetime condition effects the models’ shape discrimination accuracies. This shows that lifetime has no significant perceptual effect on network performance.

#### Varying dot density

The task here is the recognition of 3D shape in a display of reduced number of dots. Reducing the number of dots downgrades the input to be processed by low level motion systems, which in turn leads to deterioration of shape recognition performance. We compared 3 conditions in which the number of dots were 30, 60, and 90 for Helix and 45, 90, and 135 for other four shapes. We refer to these three conditions as 25%, 50% and 75% dot density display. The model shows highly accurate 3D shape recognition performance in conditions 50%, 75% and recognition of cone and sphere alone deteriorated in displays of 25% dot density as shown in Fig. 4D.

## 4 SIMULATION-2: *Recognition of point light (PL) actions using motion cues and the effect of different conditions on recognition performance*

Numerous studies have shown that people can recognize actions from sparsely disconnected points/impoverished stimuli without previous exposure, which are rarely observed in natural environments (Johansson, 1973; Troje & Westhoff, 2006). The following question is still unresolved: What visual information is crucial for rapid, accurate, and robust perception of biological movement? First class of studies (Cutting et al., 1988; Thurman & Grossman, 2008) suggest that motion computation plays a central role in biological motion perception. Studies using point-light displays provided evidence that humans may identify biological motion solely using local motion signals from joints (Johansson, 1973; Mather et al., 1992). The latter class of findings supports the hypothesis that structural or form information is also important in the visual processing of biological movement (Beintema & Lappe, 2002; Hiris, 2007; Lange & Lappe, 2006; Troje & Westhoff, 2006). In simulation-2, we investigate the contributions of structural and motion information in biological movement recognition. We trained the model with PL displays of 6 action classes (crawl, cycling, walk, jump, wave, and salute) and then tested how well the model can recognize actions in various other conditions such as inversion, scramble, oriented and masked ones. If the model exhibits a certain degree of robustness in recognizing these conditions, it would demonstrate the generalization ability of the model.

### 4.1 Stimuli Creation

Human PL action movie data are taken from dataset created by Jan Vanrie and Karl Verfaille (Vanrie & Verfaillie, 2004), and were customized according to the model requirement. We selected 6 actions: Crawl (2 views), Cycle (2 views), Walk (2 views), Jump (3 views), Wave (3 views), Salute (3 views), a total of 15 sequences, for our simulation. Each action sequence comprised 15 frames. Each frame of size 120 × 120 consists of 13 different body points (one for the head, two each for the feet, knees, hips, shoulders, elbows, and hands) each of size approximately 2 × 2 pixels were seen on a black background. Note that all the PL sequences display in place actions, as if action performing on a moving ground like a treadmill and the sample frames of each sequence is shown in Fig. 5A.

**Figure 5:**
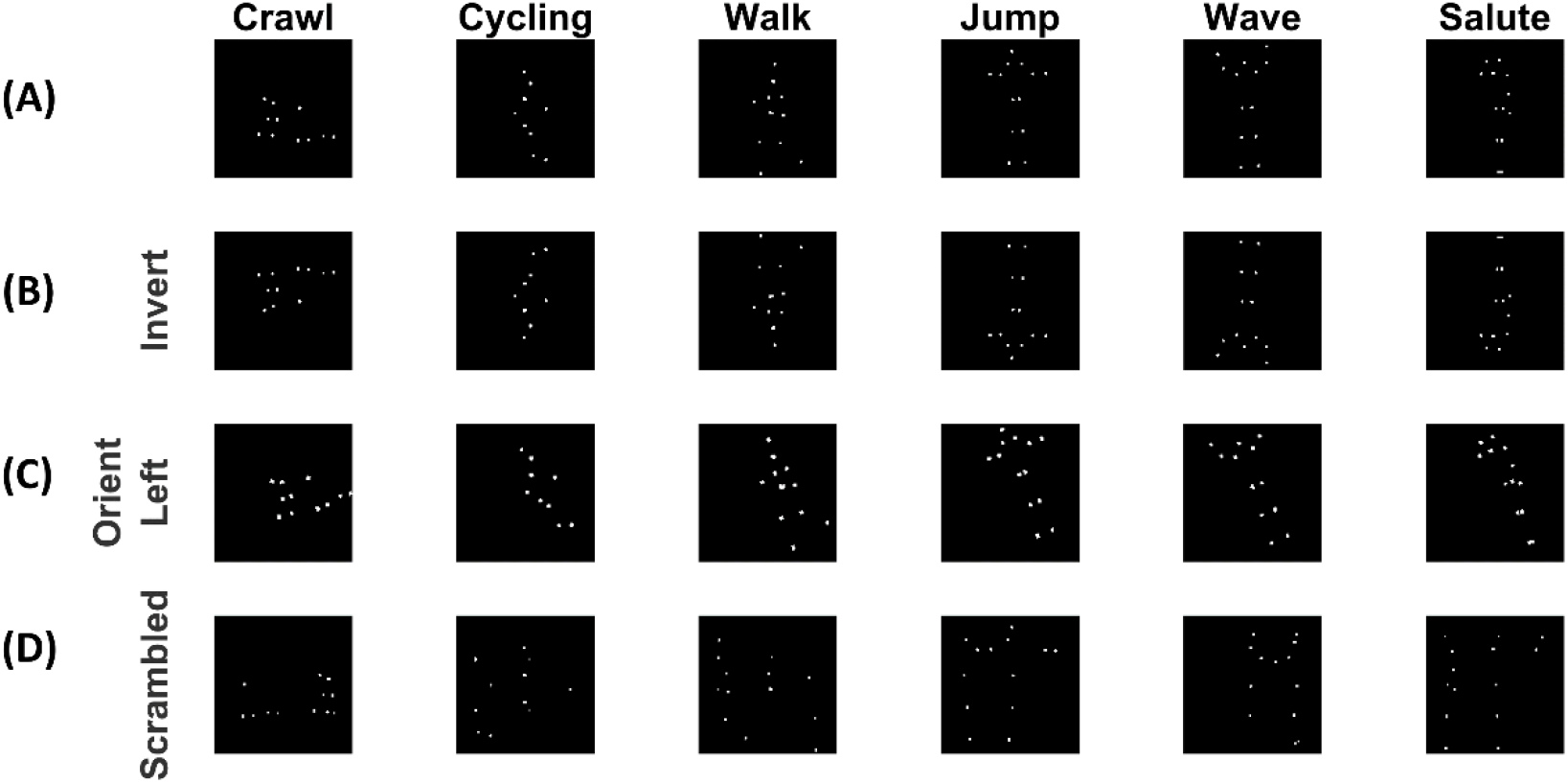
BM displays of different conditions: (A) a frame in vertical PL display (B) inverted displays, (C) displays action tilted towards lefts from vertical position, (D) displays a scrambled action.

#### Masked Stimuli

All 15 PL action sequences with different subsets of the original 13 points were created by applying 5 types of masks. Five sets of masked stimuli were composed of: (i) motion of legs (lower part of a body from hip), (ii) absence of legs (upper part of the body from hip), (iii) motion of hands and feet alone, (iv) missing motion of hands and feet, (v) missing motion of hands, feet, elbows and knees together.

#### Inverted

Each of these standard PL sequences were horizontally mirror-reflected to create upside-down versions to study inversion effects (Fig. 5B).

#### Oriented actions

Each of the vertical PL action sequences are rotated 30° in x-y plane, bidirectionally both in clockwise and anti-clockwise directions. These sequences are referred here after as 60° oriented and 120° oriented action sequences (Fig. 5C).

#### Scrambled

Scrambled motion sequences are created by shuffling the spatial configurations of each path of dots in such a way that the critical joint movements are intact as shown in Fig. 5D.

### 4.2 Training procedure

The BMNW is trained to recognize the 6 classes of PL actions. The parameters used to train CNN in BMNW is shown in Table 2. The training set contains 15 action sequences. The trained network recognition accuracy for various display formats is calculated and plotted as shown in Fig. 6. Note that training is done on undistorted stimuli and testing done on distorted (masked, inverted, oriented and scrambled) stimuli. The model performance is summarized in the following section.

**Figure 6:**
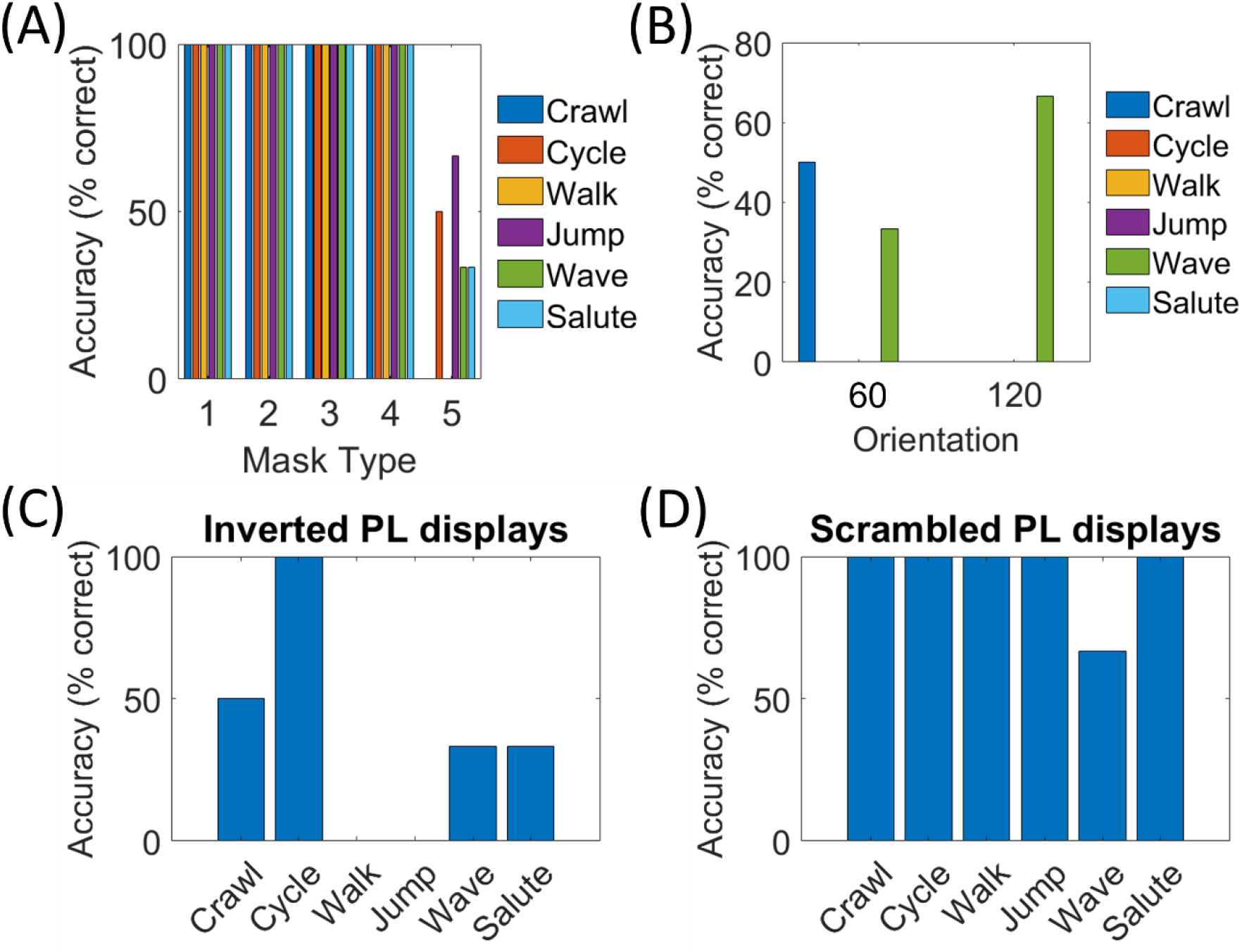
Action recognition performance of BMNW for various PL display conditions: (A) masked displays, (B) Oriented displays, (C) Inverted displays, and (D) Scrambled displays. BMNW performance is suffered severely in case of inverted and tilted actions, where as to scrambled and masked inputs performance is satisfactory.

### 4.3 Results

#### Test mask inputs

To understand the role of individual body parts in biological motion perception, masked PL displays are presented to the model. To assess the ability of the model to make use of the available information, we measured the accuracy in action recognition in each of the five mask conditions. Surprisingly, model performance (Fig. 6A) is unaffected in the first four masking conditions, indicating that the relative amount of information of either feet or hands or elbows or knees is sufficient for reliable recognition. However, the model performance has declined in the fifth mask condition where all 4 critical joint movements were absent, suggesting that joint movements in feet, hands, elbows and knees are critical and play a necessary role in recognition performance.

#### Tilt effect

To examine whether the model supports a deeper understanding of motion information in recognizing action sequences, bidirectionally rotated PL action sequences are created by changing the slope of the ground on which the action is performed. As shown in Fig. 6B for oriented inputs the model shows poor performance. Except for the actions leftward oriented crawl, wave and rightward oriented wave, all other actions are not recognized by the model.

#### Inverted actions

Inversion effect is a classic finding in biological motion perception (Pavlova & Sokolov, 2000; Proffit & Bertenthal, 1990; Sumi, 1984). It can be created by presenting PL actions upside-down. As shown in Fig. 6C, the model performance dropped significantly for all the inverted action displays except for cycling. Inverted walking and jumping actions are not recognized by the model.

#### Scrambled actions

To analyze the contribution of local dot motion trajectories in action recognition, we spatially scramble the dot motions in each action sequence. Scrambling results in a spatial dissolution of the spatial arrangement of dot pattern (Grossman et al., 2000; Kim et al., 2015). As shown in Fig. 6D, suppression of configurational form cues through scrambling has minimal effect on action recognition accuracy. The model shows high performance for all scrambled actions except for the waving action.

## 5. SIMULATION-3: *Action recognition and shape recognition using mixed stimuli where point-light actions are embedded in rotating shape*

So far, we investigated the recognition of structure from motion and biological motion with isolated stimuli. We observe that the geometrical information extracted from local motion cues plays a key role in 3D shape extraction. Similarly local motion trajectories of critical joints are important for PL action recognition. In Simulation-3 we created mixed stimuli where PL actions are embedded in a rotated 3D shape. The mixed stimuli alter the local motion trajectories of intrinsic surface dots in 3D rotating shapes and of joints in PL actions. Mixed stimuli are tested for shape and action recognition using the models SFMNW1 and BMNW trained in simulations 1 and 2.

### 5.1 Stimuli Creation and training procedure

15 PL sequences used in simulation-2 were embedded in 3 different rotating shapes: cone, cylinder, and sphere as shown in Fig. 7A. For instance, first row in Fig. 7A shows a frame from different actions embedded in a rotating cone. Note that the dots in shape and PL action are of similar texture but highlighted differently in Fig. 7A. 5 sets of each shape (5 dot positions × 15 PL actions = 75 sequences) are created by randomly rotating surface points of size 2 × 2 pixels using the methods used in simulation-1. Thus, the test set contains 225 sequences (5 sets of each shape × 3 shapes × 15 PL actions). The response of VSMN for all 225 sequences is calculated before presenting to the SFMNW1 or BMNW. Thus, when the VSMN response to a given sequence is presented to SFMNW1 the underlying shape is recognized, and the same response when presented to BMNW1, the underlying PL action is recognized. The 3D shape recognition and PL action recognition performances for mixed stimuli are estimated and the results are plotted as shown in Figs. 7B and 7C.

**Figure 7:**
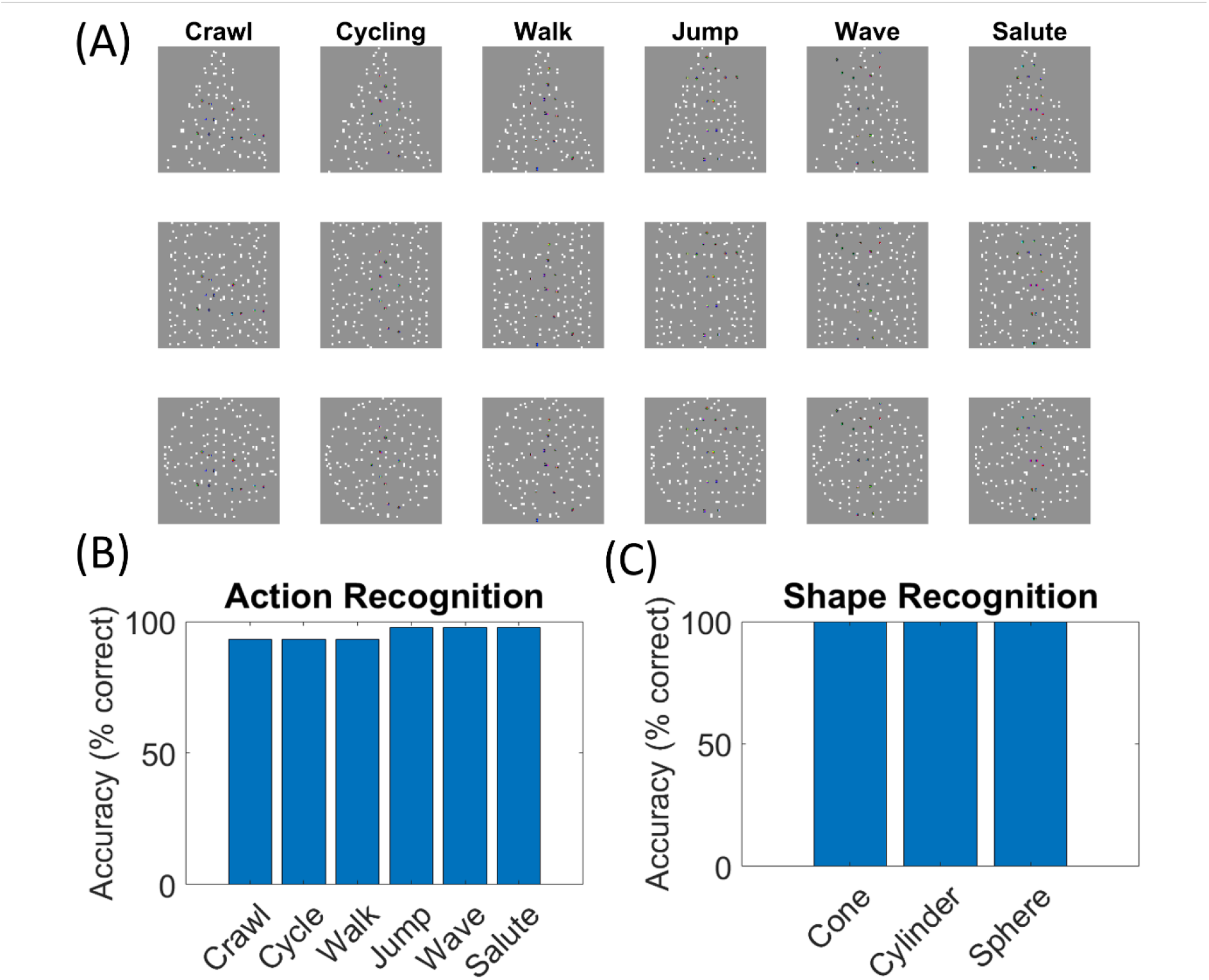
Performance of SFMNW1 and BMNW to the presentation of mixed inputs: (A) plots a frame of a sequence where PL action is embedded in a rotating 3D shape. For clarity of depiction, we differentiate dots belonging to shape and PL actions by changing pixel intensities. But in the actual stimuli, the shape dots have the intensity closer to PL action dots. (B) BMNW accuracy was lower for actions crawl, cycle and walk compared to other actions. (C) SFMNW1 showed 100% performance in shape recognition.

### 5.2 Results

SFMNW recognized all three shapes with the highest accuracy which is not surprising because the action sequence embedded within the shape has fewer dots compared to the target. Moreover, the embedded actions alter only the local motion trajectories and have minimal or no effect on the structural information. On the other hand, the BMNW performance in action recognition is surprisingly high as shown in Fig. 7B. The accuracies attained are 93.3% for actions crawl, cycle, walk and 97.8% for jump, wave and salute action classes.

## 6 SIMULATION-4: *Effect of static and dynamic noise backgrounds in 3D shape perception*

To gain insight into how the background noise (both static and dynamic) effects the shape recognition, we created visual stimuli with two overlapping random-dot patterns. The center stimulus is a rotating 3D shape defined as in simulation-1 except that each dot is of size 1 × 1 pixel, whereas the background stimulus (we call it an annulus region i.e., the region surrounding the shape which does not cover the intrinsic shape surface) is defined as one of the following 4 noise types. Note that the noise dots were present only in the annulus region surrounding the center, so that the local noise in the flow field at the central part of the shape does not lead to errors in recognition.

### 6.1 Stimuli Creation

#### Noise Type 1

The display consisting of an annular pattern of 128 randomly-placed dots of size 1 × 1 pixel. Same dot pattern is maintained throughout the sequence for every frame.

#### Noise Type 2

The display was the same as that used in the previous condition, consisting of an annular pattern of 128 randomly-positioned dots except that the random dot pattern changes for every frame.

#### Noise Type 3

This is an expanding flow condition, where all the dots move radially away from the image center point. Let *m*, *φ* be the magnitude and orientation components of dot *(x, y)*. Then the trajectory of radial motion is defined using the eqn. 6. Note that for radial trajectory magnitude (*m*) varies and orientation (*φ*) is remains constant. ‘*v*’ is the step size of dot and ‘*θ*’ takes the value zero.

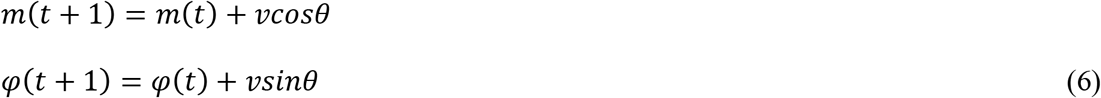

#### Noise Type 4

This is translational motion condition, where all background dots move horizontally and coherently with the same speed. The direction of background dots is the same as the direction of rotating 3D pattern. The translational trajectories are produced by updating the horizontal and vertical displacement of a dot as shown in eqn. 7.

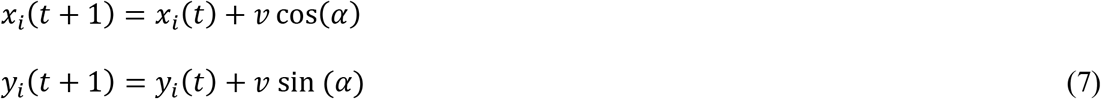

where α represents the direction of motion and the local speed is defined by ‘*v’* (= 1). α takes the value 90֯ for ellipsoid and 180֯ for all other shapes. Note that the set of dots that goes out of the square boundary will be wrapped around to reappear on the opposite side of the frame. Thus, the dot density was kept constant across the frames.

### 6.2 Training procedure

The training set consists of 300 rotating 3D shapes (5 shapes + 3 orientations (45°, 90°, 135°) + 20 random dot initial positions). Each stimulus only contained the center patch presented on a black background, without an annular surround. Here SFMNW2 is trained for shape recognition which is invariant to the orientation. Various parameters used to train CNN are given in Table 2. The trained network is used to test the stimuli with background noise.

### 6.3 Results

Here we study how the background noise influences the recognized shape of an overlapping center stimulus i.e., the rotating 3D shape. It was noted in simulation-1 that the local motion information at the boundaries is crucial for the accuracy of the derived 3-D structure rather than the ones at the intrinsic surfaces. The geometric information might be playing a key role in shape recognition. In this simulation we test whether or not the recognized 3-D structure is derived from the geometrical information. Adding the above 4 types of noises disrupts the geometrical information inherently present in each image sequence. Moreover, the dynamic background noise results in erroneous local motion information at the boundaries. Thus, we set out to test the hypothesis that the perceived 3D shape can be influenced by both static and dynamic random noise of a background stimulus, as it disrupts the geometry of a 3D shape. As shown in Fig. 8A, the static background noise has no effect on shape recognition. Except for the cylinder; for all other shapes the dynamic random motion noise reduced the accuracy of recognition (Fig. 8B). The effect is almost similar for the displays of the center patch and the background patch having same (Fig. 8D) or different motion direction (Fig. 8C).

**Figure 8:**
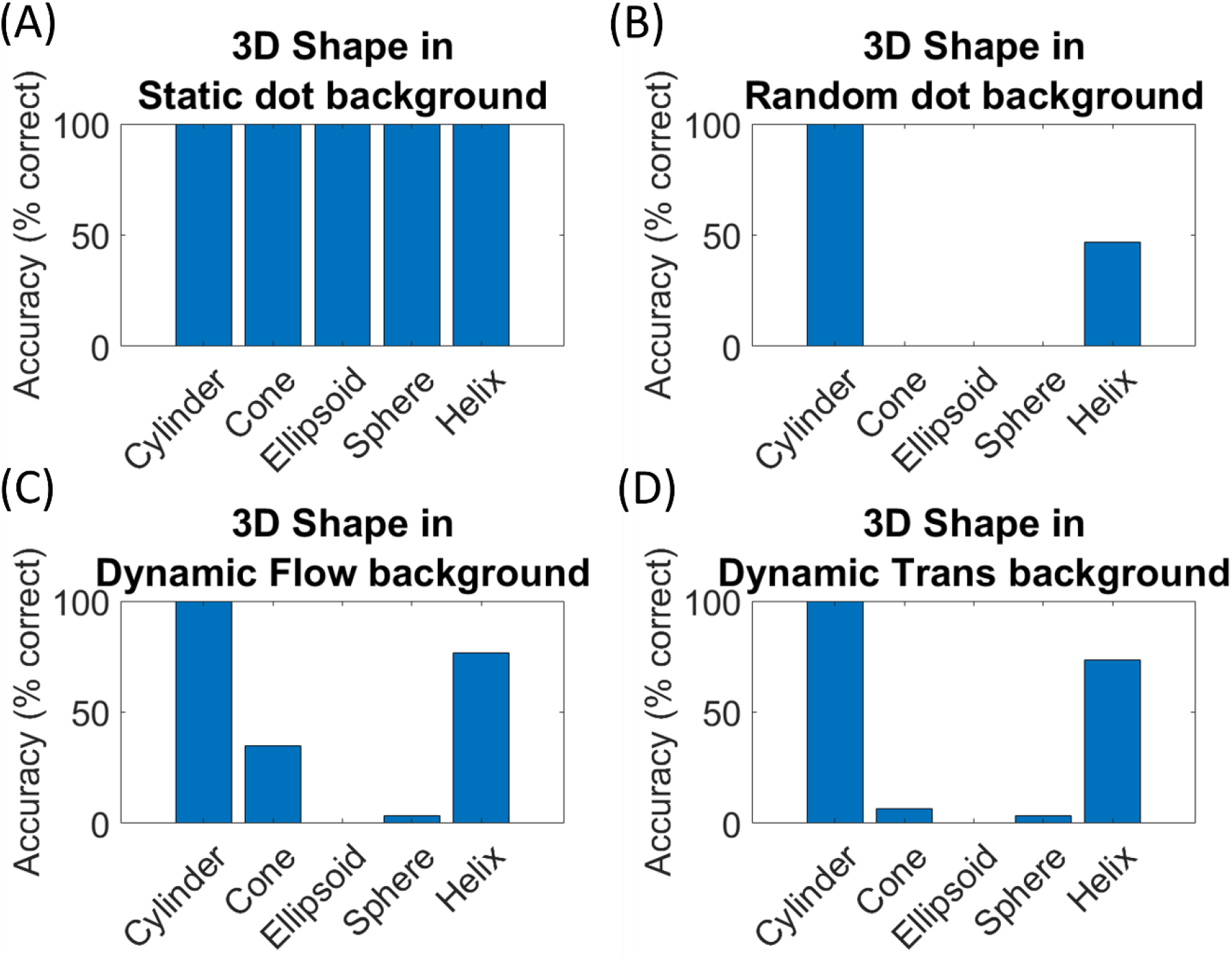
SFMNW2 performance for different noise backgrounds: Plots (A), (B), (C), and (D) represent network performance to static (fixed background dots shared by all the frames of a sequence), random (each frame has a different random backgrounds), flow (background dots move in radially outward direction), and translational (background dots move in the same direction as the dots on 3D surface) respectively. The network showed poor recognition performance to all types of dynamic dot backgrounds except for cylinder. One possible explanation could be cylinder have a straight and vertical edges whose local motion information is less disrupt by the both translational and flow noise. Similarly, cone have straight and tilted boundaries which are less effected by the flow noise.

## 7. SIMULATION-5: *Effect of static and dynamic noise backgrounds in biological motion perception*

Both experimental (Garcia & Grossman, 2008) and modeling (Giese & Poggio, 2003) studies suggest that form and motion cues together enable a reliable and robust biological motion perception (Beintema & Lappe, 2002; Casile & Giese, 2005; Hiris, 2007; Thirkettle et al., 2009; Thurman & Grossman, 2008). To analyze the relative contributions of configural form or motion-based features on the perception of biological motion, we embed the PL action sequence in a field of additional dots in four ways. This background dot motion produces a noise that negatively affects the recognition of the underlying skeleton structure.

### 7.1 Stimuli creation

First, three noise types (static, random background, outward flow) with 100 background dots were created as defined in simulation-4, except that each dot of size is maintained as 2 × 2 so that there is no distinction between the background dots and the PL action dots. In noise type 4, the noise points are placed randomly on background and each is allowed to move with the same trajectories as points from the PL action sequences present in the training set.

### 7.2 Model training

The model trained in simulation-2 on 6-category action classification is used to test the actions embedded in both static and dynamic noise.

### 7.3 Results

Simulation-2 has tested the model performance for different display formats of PL actions. In this simulation we would like to examine whether the model demonstrates robust recognition performance, as do humans, when PL actions are performed on distracting noise backgrounds. As shown in Fig. 9 the model performance is quite robust against all noise types.

**Figure 9:**
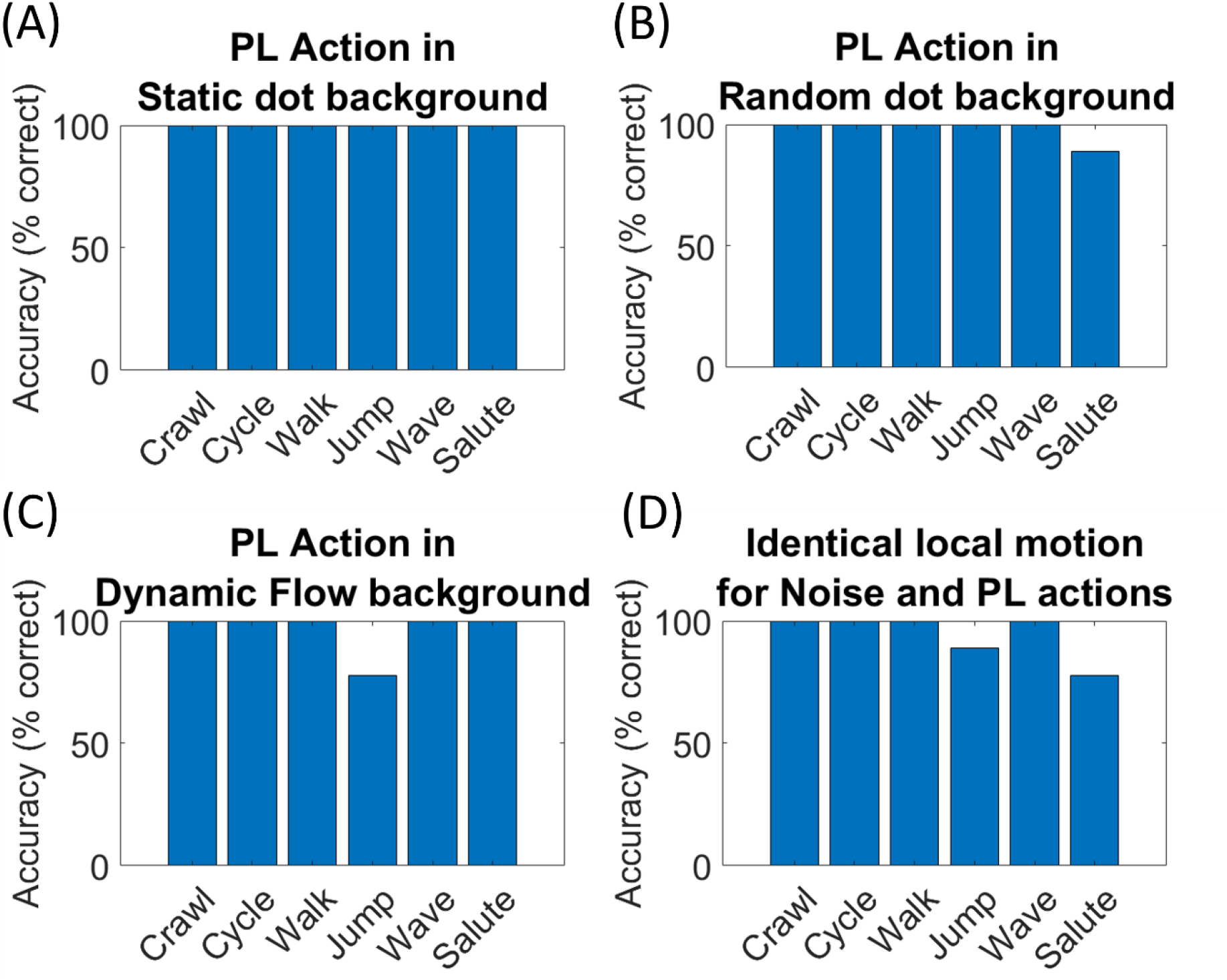
BMNW performance for different noise backgrounds: Plots (A), (B), (C), and (D) represent network performance to static (fixed background dots shared by all the frames of a sequence), random (each frame has a different random backgrounds), flow (background dots move in radially outward direction), and noise motion similar to PL action dot trajectories respectively. In all four cases network showed good recognition performance.

The performance accuracy of actions such as salute (Figure. 9B), jump (Figure. 9C) and jump in combination with salute (Fig. 9D) were affected in random dot background, dynamic flow background and identical local motion backgrounds respectively. While the static noise has no effect on model performance (Fig. 9A), the dynamic noise backgrounds showed minimum effect on action recognition.

## 8 DISCUSSION

We present a dynamic deep network model that can (i) recognize 3D shape of a rotating pattern (ii) identify an action from a PL display, under conditions of challenging noise backgrounds. The model is an expanded version of an earlier model (Gundavarapu et al., 2019). In overview, the stage-1 VSMN generates velocity-sensitive representations of moving dots. These velocity-selective population responses were used to train CNN. Two Networks SFMNW1 and BMNW were trained with standard set of sequences. The trained networks are used to test the role of various variables (motion cues) that enable the perception of 3D shape and PL action. The results of Simulation-1 are discussed below.

***(a)-*** The density of dots, when it is greater than 25%, has little or no effect on the 3D shape recognition performance. The rotating shape surface generates a 2-D flow field, in which stimulus features in the middle of the display move at a high velocity and drop to zero at the edges of the display where the dots reverse their direction and move along the opposite, hidden surface. Therefore, at the boundaries, it is the instantaneous distribution of velocities rather than the stochastic trajectories of individual dots that plays a key role in the 3D percept. But for low dot densities, the velocity distribution estimates at the object boundary are severely disrupted, which in turn reduces the models’ ability to identify shape from motion stimuli.

***(b)-*** Note that reducing individual feature stability by limiting the lifetime of individual points had no significant effect on shape identification. This may be because the information derived from individual points is preserved even after their disappearance since the overall density is preserved. VSMN generates its response by integrating information over successive frames/images. This allows features that appear separated in time to contribute to the recovery of the same object. Our results are not in agreement with earlier modeling studies (Bennett & Hoffman, 1985; Ullman, 1979, 1983), which suggest that the reducing feature stability has a large impact on the shape identification. On the other hand, our results are in agreement with the recent studies (Cheeseman et al., 2014; Norman et al., 2013) where the effect of feature lifetime is tested on discrimination of 3D shape. However, discriminating the shapes defined by the motions across two successive views (lifetime 2 condition in our simulation) lends additional support to previous experiments, where it has been found that two successive views are mathematically sufficient for the recovery of geometrical information (Koenderink & van Doorn, 1991; Todd & Bressan, 1990; Todd & Norman, 1991). Studies also have shown that subjects are quite adept at the shape identification task with displays in which dots have lifetimes of only two frames (Landy et al., 1991; Landy, Sperling, Dosher, et al., 1987). In such displays, a global optic flow field is available (although noisy), and 3D structure could, in principle, be computed from the flow field (Koenderink & van Doorn, 1987).

***(c)-*** Another important aspect of SFM concerns the recognition of oriented shapes. In our simulations 3D shape extraction is especially impaired in displays that have oriented shapes (changing the axis of rotation through angular displacement). This is because manipulations of angular displacement in the input create structural variations in 3D shape which in turn create drastic changes in VSMN responses (i.e., local patterns of velocity). These local velocity computations are the basis for the responses in stage-2 CNN. During the presentation of oriented 3D shapes, the local velocity pattern given to CNN from VSMN is very different from the patterns it was trained on. For example, in vertically upright cones, dots move from right to left (180°). When the shape has 45° orientation, the direction of motion of the dot’s changes from 180° to 225°.

However, our model could recognize 45° oriented ellipsoids with accuracy 82% and 135° oriented ellipsoid with accuracy 80%. This finding shows clearly that the model’s judgments could not have been based on a simple direct comparison/linear mapping of the two-dimensional image velocities at CNN. The higher layers of CNN are probably tuned to different variables such as global motion configurations (rotation/translation), or structural property of object (the geometrical information) that remains invariant over small variations in an object’s rotary displacement. These provide some degree of constancy over changes in an object’s angular displacement so that oriented ellipsoids are recognized correctly.

***(d)-*** We also tested the influence of motion information at boundaries and intrinsic surfaces on perception of 3D shape by introducing 5 types of masks. Occluding intrinsic surface has no effect on shape recognition, whereas masking vertical boundaries/ corners in the displays of cone, and helical curves in the displays of helix, reduces the model performance. This is because, masking them influences the shape of the extracted surface (the geometrical information) which might be playing a key role in shape extraction. Several studies (Hildreth et al., 1995; Ramachandran et al., 1988; Thompson et al., 1992) have reported that boundaries influence the 3-D interpretation of moving random-dot patterns to account for our observations.

In ***Simulation-2*** BMNW is trained to capture biological motion using 6 PL action classes. The trained network is presented with different display formats to understand the contribution of spatial and temporal features in biological motion perception. A key element in analyzing the relative contributions of structural or motion-based features is to isolate the feature under investigation by suppressing the other feature contribution. Focusing on the analysis of the contribution of local dot motion trajectories, we spatially scramble the dot motions in all 6 PL action sequences. This results in a dissolution of the spatial arrangement of the dot patterns and thus in a suppression of configurational form cues. Aiming at understanding the roles of specific stimulus features such as critical joint movements, hands or feet movement etc., we created displays of different body parts using masking technique. We also created inverted and oriented action sequences in which the underlying skeleton structure is different from the one in the training set. Aspects of the results from Simulation-2 are discussed below.

***(a)-*** The model performance has significantly dropped when inverted action sequences are shown. The model results are inconsistent with human judgements in various direction discrimination tasks (Bardi et al., 2014; Blake & Shiffrar, 2007; Reed et al., 2003; Troje & Westhoff, 2006).

***(b)-*** On similar lines, while presenting oriented stimuli, model judgments show that the actions were judged correctly only when the figure’s orientation is consistent with orientation of the training stimuli and a substantial drop in accuracy was seen for differently oriented PL actions.

***(c)-*** On the other hand, the model showed better accuracy in recognizing actions even when dot configuration is disrupted by scrambling, where critical joint movements are intact. Model results are in agreement with the studies reported by Troje and Westhoff (Troje & Westhoff, 2006).

These results indicate that the local motion signals of joint movements are an important cue in biological motion perception. Neurons in the visual system tuned to such critical joint movement information are described as “life detectors” by Troj & Westhoff (Troje & Westhoff, 2006a). Particularly, motion of the feet has been proposed as the primary signal being used by such a life-detector mechanism. Empirical studies (Mather et al., 1992) on identifying the most important parts of the body, supports this idea that feet and hands exclusively can result in accurate direction discrimination of PL walkers. Our findings in masked displays extend the results of several of these studies in exploring role of parts and wholes on the perception of biological motion.

***(d)-*** We presented masked PL actions to identify the relative importance of different body parts. The model accuracy was highest in the first four mask conditions. The third mask condition reflects a genuine processing advantage for the motion of hands and feet. In the fourth mask condition, where hands and feet are missing, the model performance was still accurate suggesting that the movement of knees and elbows contribute significantly to the action recognition. In the fifth mask condition, where all the four parts - hands, feet, elbows and knees - are missing together, the model performance dropped significantly. This suggests that the local motion information provided by critical joints such as elbows, knees, hands and feet might carry critical discriminative information which resulted in accurate PL action recognition.

At this point we would like to draw comparisons of the model with various cortical regions involved in biological motion perception. Superior Temporal Sulcus (STS) has often been implied undisputedly in the perception of biological motion in imaging studies (Beauchamp et al., 2002; Bonda et al., 1996; Grossman et al., 2000; Puce et al., 1998; Thompson et al., 2005; Vaina et al., 2001), electrophysiological investigations (Oram & Perrett, 1994, 1996), and lesion studies (Cowey & Vaina, 2000; Vaina & Gross, 2004). Local motion signals are believed to be processed in middle temporal (MT) gyrus (Ptito et al., 2003; Vaina et al., 2001) and the kinetic occipital area (KO) (Santi et al., 2003; Vaina et al., 2001). Studies (Downing et al., 2001; Grossman et al., 2000) have reported that activation in these areas is similar to both control and scrambled stimuli (stimuli with identical motion signals but altered spatial arrangement). As the signals from MT and KO feed into the STS, local motion signals might contribute to the biological motion recognition in STS.

In Simulation-3, we adopted the models (SFMNW1 and BMNW) trained in simulations 1 and 2 to test the recognition performances on mixed stimuli where PL actions are embedded in 3D rotating shapes. Both networks reported highest recognition accuracies. Predominantly the high discrimination performance of the BMNW for action recognition showed that, even when shape dots in mixed stimuli alter the local motion trajectories of the embedded action dots, the global motion structure of the stimulus is preserved. Global motion can theoretically be derived from integration of local motion signals, or the trajectories of the point-lights over time (Giese & Poggio, 2003). The global motion signals may help to segment the stimulus from the background. Thus, extraction of the embedded action is in fact facilitated by enabling figure ground segregation. Our results are consistent with the results by Neri et al. (Neri et al., 1998) where they reported a remarkable efficiency of human observers in the temporal integration of biological motion in noise.

In ***Simulation-4,*** our goal is to evaluate 3D shape recognition performance as a function of the background noise type. We trained SFMNW2 that recognizes shape irrespective of its orientation. The network performance is tested by presenting inputs where shapes are embedded in various noise backgrounds. The simulation results show that the motion noise at rotating surface boundaries has a significant impact on 3D shape recognition and suggest that the motions of high curvature regions (boundaries in this case) and corners provides a strong cue to the recognition of 3D shape. One possible neural explanation of our results is that the invariant motion information in V1 is achieved by the activity of motion sensitive end-stopped neurons (Pack et al., 2003; Pack & Born, 2001). These neurons provide both the motion and form signals that are unambiguous with respect to the aperture problem. Studies have shown that structure-from-motion is a highly interactive process that incorporates not only motion cues but also form cues (Kourtzi et al., 2008). In neurobiological terms, SFM is supported by a cascade of brain regions spanning dorsal and ventral streams, such as MT/V5, V3A, and LOC (Brouwer & van Ee, 2007; Orban et al., 1999; Paradis et al., 2000; Raemaekers et al., 2009; Vanduffel et al., 2002). We believe that background noise invariant 3D shape recognition from motion-defined surfaces can be achieved by incorporating form cues.

In ***Simulation-5*** we tested the sensitivity of proposed BMNW to noise in the PL action displays. Simulations showed that the perception of PL actions is robust against background noise. The results are consistent with earlier studies (Beintema & Lappe, 2002) which showed that humans can recognize actions in PL displays where local motion signals are eliminated, as long as stimuli preserve a certain degree of the global form. The contributions of form and motion to the perception of PL displays are subject to controversy in the discussion. While some studies claim that local motion signals are critical, others emphasize the role of global form cues. Our results indicate that together local motion signals and global form cues were used in combination for the generalized action recognition performance. Even though the background noise disrupts the skeletal structure of PL action sequences, integration of local motion information over time (in the form of VSMN response in our model) mediates the salience of form pattern. The body shape information learned by CNN in BMNW (some authors refer to it as *configural form cues* (Bardi et al., 2011; Lange & Lappe, 2006; Thompson et al., 2005) will extract the embedded action from the noise background.

Thus, we believe that PL action recognition is a two-step process. The local motion information and the form built up by integrating these local motion cues (configural form) seems to be sufficient for biological motion perception, and perhaps the explicit form information is not critical for biological motion recognition. Current models (Giese & Poggio, 2003) of biological motion processing suggest that human form and motion information are initially processed through separate pathways, then integrated in action perception, which is contrasted with the current model. Our modeling approach is actually in line with the recently proposed two-process theory of biological motion processing (Hirai & Senju, 2020). On the other hand, early computational models of SFM have an extreme sensitivity to noise in the visual input and suggested various ways to overcome this sensitivity (Adiv, 1985; Landy, Sperling, Perkins, et al., 1987; Ullman, 1984; Waxman & Wohn, 1988).

All the observations reported in this paper are based on experimental studies involving laboratory stimuli such as random dots and PL actions. Although PL displays have served as useful research tools, they have limitations. The PL displays provide incomplete structural information about the human body. For these reasons, the observers’ judgments in various studies are questionable and it is difficult to confirm that the identification of biological movement is carried out from motion cues alone. Also, the rarity of these laboratory stimuli in natural scenes raises a question about the generalizability of experimental findings. Future models of SFM perception and biological motion perception should be trained and tested on natural stimuli with more ecological considerations.

### Pros and cons of the current model

One attractive feature of the proposed model is that it provides a simple biologically plausible architecture for solving problems of SFM computation and biological motion perception. Many studies emphasized that both spatial and temporal pathways contribute to the generalization of SFM and biological motion perception. However, the 5 simulation studies described in this paper indicate that separate mechanisms may seem to underlie these perceptual phenomena. SFM is related to rotation in rigid depth stimuli, whereas biological motion perception is related to nonrigid motion stimuli that carries form and temporal sequence information.

Initially we hypothesized that the unified architecture will suffice to understand the mechanisms used by these two perceptual phenomena. The model showed more general, robust biological motion recognition and SFM recognition (in noiseless displays). However, with the current results on noise SFM displays, it could not be explained that the low-level motion features which further mediate the form cues, lead to 3D shape recognition in rotating surfaces. It would seem that SFM perception involves mechanisms that use more elaborate form or motion information that help to recognize and segment the pattern of movement (Aggarwal & Cai, 1999) in ambiguous displays. Recent priming studies (Pastukhov et al., 2013) reported that the sensory memory of SFM is shape specific, involves higher-level representations of object surfaces and shapes.

On the other hand, biological motion perception involves more specialized and unique evolutionary mechanisms (Neri et al., 1998). Neural responses of the anterior superior temporal polysensory area of the macaque temporal cortex is said to be sensitive to the human form (Oram & Perrett, 1994). On the other hand, SFM is a highly interactive process that incorporates not only motion cues but also form cues. The claim of distinct mechanisms for SFM and biological motion has been investigated before (Neri et al., 1998; Poom & Olsson, 2002) where they compared spatiotemporal integration characteristics of three different moving stimuli: biological motion, translation, rotation in depth, by plotting spatiotemporal summation curves. They found these curves appear to be different for each of the above three stimuli, thus suggesting that 3 different mechanisms exist for these three types of moving stimuli.

The next natural step in improving the model would be to introduce a second order motion pathway. Such architecture may prove to be a more generic model of motion perception, capable of solving a variety of perceptual problems using natural stimuli, which would be difficult to solve using current architectures.

### Relation with other models

Finally, we discuss the relationship between our model and other models of SFM perception and biological motion perception. Many modeling studies support the idea of interactions and the functionality segregation among two parallel motion and form pathways. In particular, the modeling approach of Giese and Poggio (Giese & Poggio, 2003) suggests that the visual input is processed largely independently over a hierarchy of stages in two parallel pathways thereby incorporating filtering and integration mechanisms. In line with this interpretation, the model by Thurman et al. (Thurman et al., 2010) included separate form and motion pathways. A hierarchical model of point-light walker (PLW) proposed by Troje (Troje, 2008, 2013) consists of at least two interacting modules processing both local motion and global SFM information. Another computational model of PLW by Lappe (Lange et al., 2006) has two successive stages. First stage of the model consists of template cells for whole-body postures, which calculate the stimulus orientation. In the subsequent second stage, temporal order is calculated that identifies the global form. Output of second stage is used to make decision on the global motion aspects of the stimulus. A neural architecture consisting of a two-stream convolutional neural network, proposed recently by Peng (2021,) supports the two stream hypotheses for biological motion perception. The model consists of a spatial CNN to process appearance information, a temporal CNN, to process optical flow information, and a fusion network to integrate the features extracted by the two CNNs and make final decisions about action recognition.

The model proposed in this paper has two successive stages in which local motion information and global form information are extracted sequentially. The proposed model is similar to the two successive stage architecture proposed by Lappe (Lange et al., 2006). However, the neural responses in each stage are very different. More details about the feature maps that are tuned to optic flow information in the input stimuli were described in our companion paper (Gundavarapu and Chakravarthy, 2022). A similar concept of contribution of neurons tuned to optic flow information to biological motion can be seen in (Peng et al., 2021).

## 9. METHODS

### Piecewise linear sigmoid function

The piecewise linear sigmoid activation function is an efficient approximation of the logistic function and implements the essential thresholding and saturation behaviour more quickly than a smooth logistic function. Here the neuron responds only if the input is as large as the threshold *θ_l_*, and saturates at *θ_u_*. The output activation values are limited to [0 .. 1]. In our simulations *θ_l_* and *θ_u_* are not modeled as a free parameter instead they were chosen from the instantaneous activities of NF neurons. Let *η(s)* be the activity produced on NF at settling time step *s*. *η_ij_* is the activity of the NF neuron *(i, j)*. We considered maximum activity produced in NF as the upper bound and the difference between maximum and minimum activities as a lower bound. Note that nonlinearity is calculated for each step in the settling process. This modification helps in producing a single-winner neighborhood.

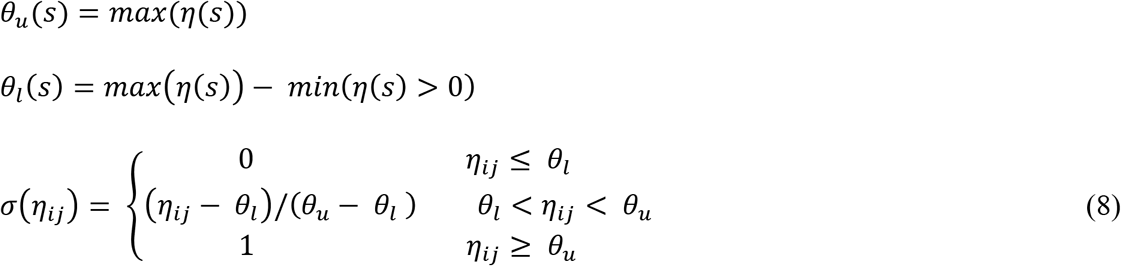

#### Creating rotating shapes

Five rotating shapes are created using the following procedure.

##### Sphere

The sphere is rotated smoothly in the image plane using eqn. (9).

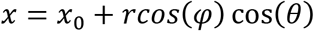

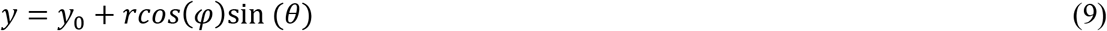

where *0 <= θ <= 2π, -π/2 <= φ <= π/2*. *r* is radius of a sphere centred at *(x_0_, y_0_)*. With geographical intuition *θ* can be referred as lines of longitude and *φ* can be referred as lines of latitude.

##### Ellipsoid

An ellipsoid squashed along each axis by a, b is defined parametrically as

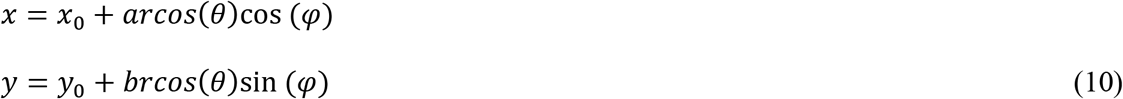

where *0 <= θ <= 2π, -π/2 <= φ <= π/2*. *r* is radius, *(x_0_, y_0_)* is centre, and *a, b* are constants taken as 0.5, 50.

##### Cylinder / Cone

Let v1, v2 be two random variables distributed uniformly on the interval [0,1] that generate the latitude β1 and longitude β2 as follows

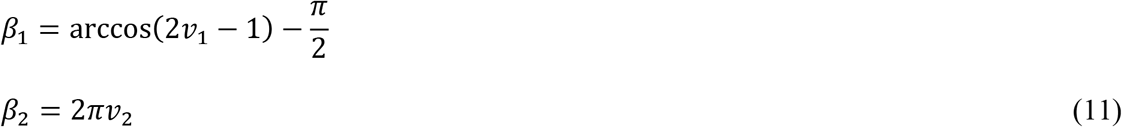

Then the coordinates for cylinder or cone can be computed as follows. Cone is a special case of cylinder with one of the facet radii is equal to one.

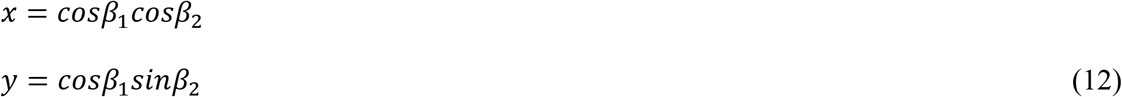

##### Helix

Helical motion is produced when one component of velocity is constant in magnitude and direction while the other component is constant in speed but uniformly varies in direction.

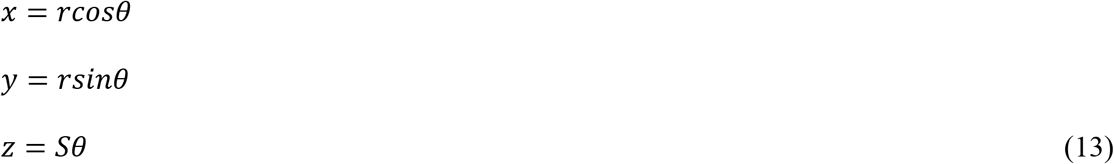

Where *r* is the radius of helix, *θ* is the winding angle and *S* is the distance between the turns.

## AUTHOR CONTRIBUTIONS

AG performed designing, coding, running simulations and manuscript preparation. VC performed designing the model and manuscript preparation.

